# Microtubule stabilization drives 3D centrosome migration to initiate primary ciliogenesis

**DOI:** 10.1101/080556

**Authors:** Amandine Pitaval, Fabrice Senger, Gaëlle Letort, Xavier Gidrol, Laurent Guyon, James Sillibourne, Manuel Théry

**Author notes:** Current address: Autolus Limited, Forest House, 58 Wood Lane, London, W12 7RP, England (James Sillibourne). Correspondence to James Sillibourne; or Manuel Théry.

## Abstract

Primary cilia are sensory organelles located at the cell surface. Their assembly is primed by centrosome migration to the apical surface. Yet surprisingly little is known about this initiating step. To gain insight into the mechanisms driving centrosome migration, we exploited the reproducibility of cell architecture on adhesive micropatterns to investigate the cytoskeletal remodeling supporting it. Microtubule network densification and bundling, with the transient formation of an array of cold-stable microtubules, and actin cytoskeleton asymmetric contraction participated in concert to destabilize basal centrosome position and drive apical centrosome migration. The distal appendage protein Cep164 appeared to be a key actor involved in the cytoskeleton remodeling and centrosome migration, whereas IFT88's role seemed to be restricted to axoneme elongation. Together our data elucidate the hitherto unexplored mechanism of centrosome migration and show that it is driven by the increase and clustering of mechanical forces to push the centrosome toward the cell apical pole.

## Introduction

The centrosome, the major microtubule-organizing center of the cell, can transform to a primary cilium, a sensory organelle protruding from the cell surface, when the cell exits the cell cycle and enters a quiescent state (Bornens, 2012). Sensory function is endowed to the primary cilium by transmembrane receptors, which localize to and concentrate within the extracellular part of the primary cilium (Nigg and Raff, 2009). As a sensory organelle, the primary cilium plays important roles in embryonic development and cellular homeostasis. Genetic mutations that result in the failure to form a primary cilium cause developmental defects including polydactyly, craniofacial defects and heart malformation, and highlight the crucial role of this organelle in development (Goetz and Anderson, 2010).

Primary cilium assembly is a complex and highly coordinated process, which is reflected by the large number of primary ciliogenesis effectors and their diverse functions (Kim et al., 2010; Wheway et al., 2015). This multi-step process begins with the assembly of a ciliary vesicle at the distal end of the mother centriole (Knödler et al., 2010; Nachury et al., 2007; Westlake et al., 2011; Lu et al., 2015). Recent work has shown that Cep164, a distal appendage protein (Graser et al., 2007), plays an important role in anchoring factors regulating centriole elongation (Cajánek and Nigg, 2014) and the formation of the primary ciliary vesicle (Schmidt et al., 2012). Following ciliary vesicle formation, tubulin dimers are added to the minus ends of the centriolar microtubules of the mother centriole to form an axoneme, which protrudes from the cell surface and is ensheathed by ciliary membrane. This is dependent upon the activity of a multi-subunit complex known as the intraflagellar transport (IFT) complex interacting with kinesin and dynein molecular motors (Lechtreck, 2015). Finally, basal body anchoring to the cortex is mediated by the mother centriole's distal appendages and several of their components have been identified. Cep83 is a key distal appendage protein (Joo et al., 2013) that is responsible for anchoring four other components, Cep89 (Cep123/CCDC123) (Sillibourne et al., 2013), SCLT1, Cep164 (Graser et al., 2007) and FBF1 to the distal appendages (Tanos et al., 2013). During, or after, the process of ciliary vesicle formation and axoneme extension, the mother centriole migrates to the cell surface, where it attaches to the cortex (Singla et al., 2010; Reiter et al., 2012). Despite much information regarding basal body maturation and anchoring, and the players involved in the regulation of centrosome positioning (Barker et al., 2015), the physical mechanism powering centrosome displacements and migration to cell apical pole is poorly understood.

Microtubules regulate centrosome positioning at the cell center by exerting pushing and pulling forces (Burakov et al., 2003; Zhu et al., 2010; Kimura and Kimura, 2011). They have also been shown to support centrosome migration away from cell center toward the cell surface by the production of pulling forces during immune synapse formation (Yi et al., 2013) or mitotic spindle positioning (Morin and Bellaïche, 2011). Interestingly, recent numerical simulations suggested that asymmetric pushing forces could also efficiently promote centrosome off-centering (Letort et al., 2016). Whether pushing forces on the basal pole and/or pulling forces from the apical pole are involved in centrosome migration during primary ciliogenesis remains to be uncovered.

Actin cytoskeleton is also involved in the regulation of centrosome positioning and ciliogenesis (Pitaval et al., 2010; Dawe et al., 2009; Kim et al., 2010). Acto-myosin contractility is required for basal body migration to cell apical pole (Hong et al., 2015; Pitaval et al., 2010) but its hyper-activation impairs primary cilium formation and cilium elongation (Rao et al., 2014; Pitaval et al., 2010; Kim et al., 2010). In parallel, disruption of actin filament formation, either by depletion of Arp2/3 or treatment with a low dose of cytochalasin D, promotes primary ciliogenesis (Sharma et al., 2011; Kim et al., 2010). Indeed, branched actin filament formation impairs the recruitment of primary ciliogenesis effectors to a region surrounding the centrosome referred to as the pericentrosomal preciliary compartment (Kim et al., 2010; Rao et al., 2014). Recent work has demonstrated that the Arp2/3 complex is present at the centrosome where it promotes the nucleation of actin filaments (Farina et al., 2016). This centrosomal network needs to be disassembled for the centrosome to detach from the nucleus and move to the periphery during immune synapse assembly (Obino et al., 2016). A similar regulation of the centrosome-nucleus link has been proposed to be required for centrosome migration during ciliogenesis (Adams et al., 2012; Dawe et al., 2009) although the physical mechanism powering centrosome motion has not yet been established. Thus, the exact mechanism by which actin network architecture and contractility regulate centrosome migration during ciliogenesis remains unclear and needs further characterization.

In this paper, we exploited the reproducibility of cell architecture on adhesive micropattern to investigate the mechanisms driving apical centrosome migration during primary ciliogenesis. Dramatic remodeling of the actin and microtubule cytoskeletons was found to drive apical centrosome movement and relied on the activity of molecular motors. A candidate-based siRNA screen of primary ciliogenesis effectors identified a role for the distal appendage protein Cep164 in centrosome movement. These data characterize in detail the previously understudied process of centrosome migration and identify unreported roles of known ciliogenesis effectors in this process.

## Results

### Centrosome migration in serum-starved cells

Previous studies have shown that the culture of cells on adhesive micropatterns promotes reproducible organelle positioning and results in a defined intracellular architecture (Théry et al., 2006; Pitaval et al., 2013). We have shown that primary ciliogenesis can be induced in isolated single retinal pigment epithelial 1 (RPE1) cells cultured on disc-shaped micropatterns and is influenced by cell confinement and contractility (Pitaval et al., 2010) as it is the case in vivo (Blitzer et al., 2011). Highly confined cells exhibiting low contractility form primary cilia more frequently than less constrained cells exhibiting higher contractility. For this reason, we chose to monitor centrosome migration during primary ciliogenesis in RPE1 cells growing on small disc-shaped micropatterns with an area of 700 μm2. Furthermore, the increased cell height associated to cell confinement offered the possibility to monitor basal body migration over a few microns (Figure 1A, B). Thereby we could distinguish centrosome migration and axonemal elongation defects in cells where ciliogenesis was impaired.

**Figure 1.**
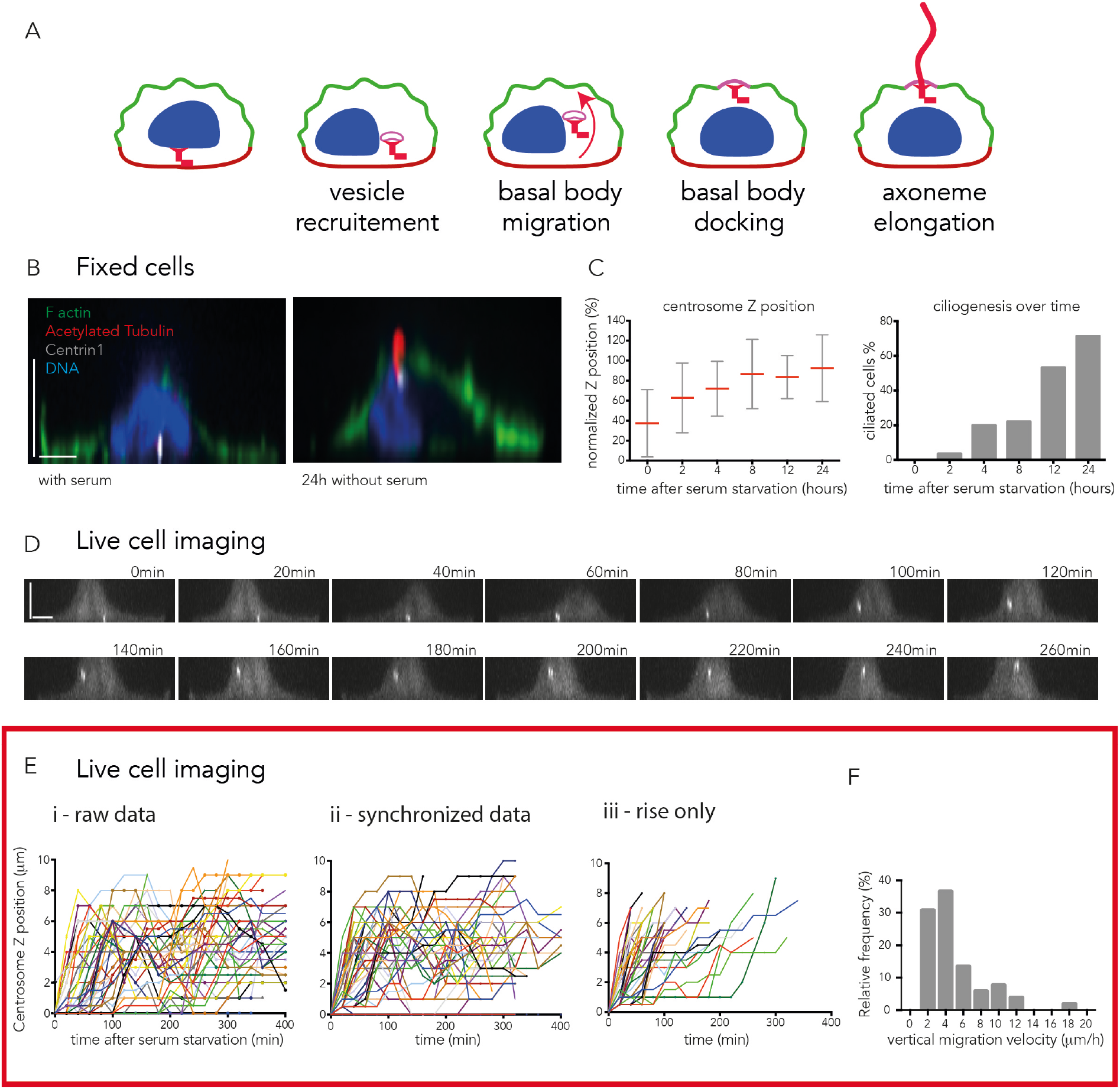
Adhesive micropatterns facilitate the study of centrosome migration during primary ciliogenesis. **A)** Primary ciliogenesis is a multi-step process that is proposed to begin with centrosome maturation and the formation of a ciliary vesicle at the distal end of the mother centriole, after which the centrosome migrates to the apical surface and attaches to the cortex. Full extension of the axoneme occurs once the mother centriole is anchored to the cortex. **B)** Micropatterned RPE1 cells expressing EGFP-centrin1 (white), cultured in the presence or absence of serum for 24 hours, stained with phalloidin (green), to visualize F-actin, acetylated tubulin antibody (red) to label the cilium and DAPI to stain the DNA (blue). Scale bars represent 5 microns. **C)** Measurement of centrosome Z position, as a percentage of nuclear height. Migration started within 2 hours of serum starvation and appeared completed 6 hours later. Measurement of the proportion of ciliated cells showed a delayed process compared to centrosome migration (one experiment, n=60 cells per condition). **D)** Side view of a representative time lapse imaging of serum-starved RPE1 EGFP-centrin1 cells on micropatterns. Centrosome migration was engaged 2 hours after starvation and completed 2 hours later. Scale bars represent 5 microns. **E)** Representation of time lapse centrosomes movement in serum-starved RPE1 EGFP-centrin1 (data of 3 independent experiments, n=53 cells). (i) The graph represents all the raw data. (ii) The centrosomes trajectories are synchronized ie they start when the centrosome leaves the basal pole. (iii) The plotting of the centrosomes trajectories is limited to the maximal z position ie they come to an end when centrosomes reach apical pole. **F)** Frequency distribution of centrosome migration velocity from basal pole to apical pole.

RPE1 cells expressing EGFP-centrin1 were cultured either in the presence or absence of serum for 24 hours and were then fixed and stained with phalloidin to label F-actin, DAPI to label DNA and an antibody to acetylated tubulin to label cilia. This revealed that the centrosome was located at the basal surface in cells cultured in the presence of serum, while those cultured in the absence of serum had formed cilia and the centrosome was at the apical surface (Figure 1B). To determine the timing of centrosome migration, RPE1 cells expressing EGFP-centrin1 were serum-starved over a 24 hours period, stained with an acetylated tubulin antibody and the axial position of the centrosome was determined and expressed as a percentage of nuclear height. Ciliated cells were also enumerated. Surprisingly, centrosome migration was found to occur as soon as 2 hours after serum starvation and by 8 hours the centrosome was located at the apical surface (Figure 1C). As expected, primary cilium formation took longer to complete with the maximal number of ciliated cells only being reached after 24 hours of serum (Figure 1C). Monitoring of centrosome movement in serum-starved RPE1 EGFP-centrin1 cells by live imaging provided a more detailed picture of the 3D migration mechanism in individual cells (Figure 1D). The ignition of centrosome take off was preceded by a lag period. This lag period varied from few minutes to few hours. These variations blurred the analysis of centrosome motion trajectories (Figure 1E-i). A clearer picture could be obtained by synchronizing trajectories at the very moment of centrosome take off (Figure 1E-ii). However centrosome wandering after the vertical migration step maintained some confusion. We then limited the plotting of the trajectory to the elevation step and could thus specifically analyze it quantitatively (Figure 1E-iii). We found that the duration of centrosome motion from basal to apical pole was quite variable, ranging from 30 to 300 minutes. The corresponding centrosome velocity was comprised between 2 and 20 µm/min with a median value close to 5 µm/min (Figure 1F). Together, these data demonstrated that centrosome migration was one of the early steps of primary ciliogenesis that occurred rapidly after serum starvation, and that micropatterns enabled the process to be easily monitored and quantified. Interestingly, the vertical displacements from the basal to the apical pole was about a hundred time slower than the reported velocities for centrosome migration toward immune synapse in T lymphocytes (Yi et al., 2013), or sperm aster centration in sea urchin eggs (Tanimoto et al., 2016), suggesting that the positioning mechanism was likely to differ from the pulling forces involved in these two examples.

### Specific implication of ciliogenesis effectors in centrosome migration or axoneme elongation

Next, we decided to investigate the role of known primary ciliogenesis effectors in centrosome migration by using siRNA to mediate their depletion. Candidates were chosen to reflect the diversity of ciliogenesis effectors and included Cep164 (Graser et al., 2007), Cep123 (Cep89/CCDC123) (Sillibourne et al., 2013; Tanos et al., 2013), intraflagellar transport 20 (IFT20) (Follit et al., 2006), partitioning defective 3 (Pard3) (Sfakianos et al., 2007), nesprin2, meckelin (Dawe et al., 2009), pericentrin (Jurczyk et al., 2004), Kif3A (Lin et al., 2003), IFT88 (Pazour et al., 2000). The role of kinesin light chain (KLC1) and emerin in the regulation of centrosome anchoring to the nucleus were also tested (Schneider et al., 2011; Salpingidou et al., 2007; Roux et al., 2009). Small-interfering RNA-mediated protein depletion was used to assess the role of each candidate in centrosome migration (Supplementary Figure S1). RPE1 cells previously treated with siRNA were plated onto micropatterns and after 24 hours of serum starvation. The deletion of all tested proteins except pericentrin and IFT88 had significant deleterious effect on centrosome migration suggesting these two later were specifically involved in axoneme elongation (Figure 2A). The finding that Kif3A depletion impact centrosome migration agrees with previously published data showing that defective ciliogenesis was associated to basal bodies mispositioning in mouse hair cells (Sipe and Lu, 2011) and zebrafish photoreceptors (Pooranachandran and Malicki, 2016). The effect of IFT20, meckelin and nesprin2 depletion on basal body positioning confirmed earlier observations in mouse kidney (Jonassen et al., 2008) and in cultured kidney cells (Dawe et al., 2009). To corroborate our results with those of others, we determined the frequency of primary cilium formation and found that it was significantly reduced after treatment with siRNA targeting the candidate mRNAs (Figure S2A). In addition, cilium length was measured and found to be reduced compared to the control siRNA, except where the cells were treated with siRNA to meckelin, KLC1 or Ift20 (Figure S2B). Together these results suggested that depletion of the candidate proteins was successful.

**Figure 2.**
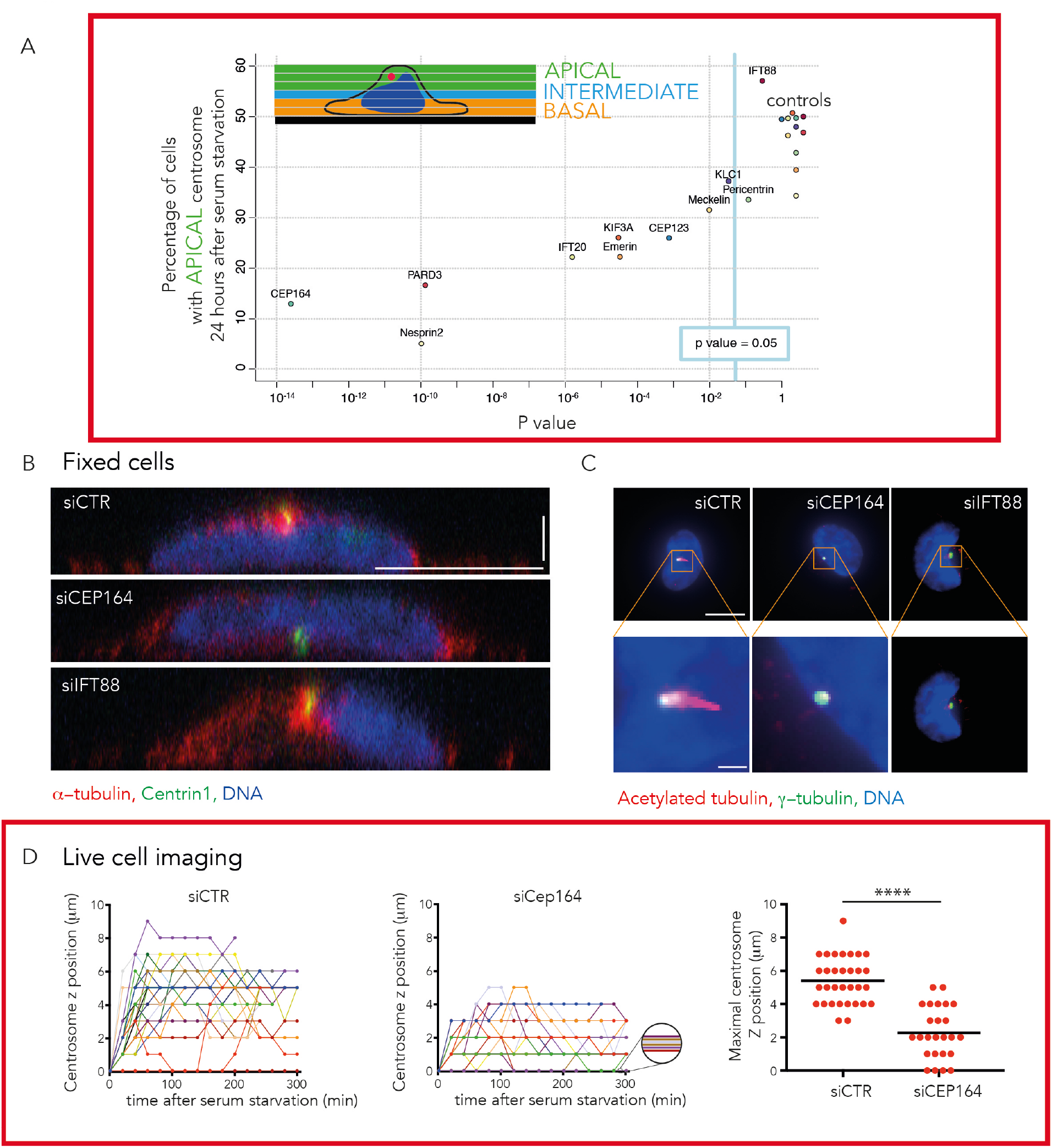
Implication of known ciliogenesis effectors in centrosome migration. **A)** RPE1 EGFP-centrin1 cells were treated with siRNAs targeting known primary ciliogenesis effectors for 24 hours to investigate their potential role in centrosome migration during primary cilium formation. The proportion of cells displaying a centrosome located more than 3 μm above the basal pole was determined and normalized to that of the non-targeting control siRNA for each condition. See Supplementary Figure S1 for the siRNA sequences and Supplementary Figures S2A-B for the effect on the rate of ciliated cells and the length of primary cilia. **B)** Side views of serum-starved RPE1 EGFP-centrin1 (green) cells stained with DAPI to label the DNA (blue) and an antibody to α-tubulin (red) to stain microtubules. XY scale bars represent 10 µm and Z scale bar represents 2,5 µm. **C)** Staining of RPE1 cells with DAPI (blue) and with antibodies to acetylated tubulin (red), γ-tubulin (green). Images show maximal projection of Z stacks. Scale bars in zoomed images represent 1 µm. **D)** Representation of synchronized time lapse centrosomes movement in serum-starved RPE1 EGFP-centrin1 treated with non-targeting control siRNA (1 experiment, n=32 cells) or siRNA against siCep164 (1 experiment, n=25 cells) (left and middle). The graph represents the maximal centrosome z position for cells treated with non-targeting control siRNA and with siRNA against siCep164 (right).

Two candidates, Cep164 and IFT88, were selected for further analysis as their depletion (Supplementary Figure S2C) resulted in opposing phenotypes, with depletion of Cep164 blocking centrosome migration and IFT88 ablation having no effect. Confocal imaging and side view reconstructions confirmed our initial results and showed that after serum starvation the centrosome was at the basal surface in Cep164-depleted cells, while in Ift88-depleted cells it was at the apical surface (Figure 2B), but neither possessed a cilium, in contrast to control siRNA-treated cells (Figure 2C). Cep164 is a core component of mother centriole distal appendages (Graser et al., 2007). It is involved in the docking of the ciliary vesicle (Schmidt et al., 2012). More generally, distal appendages are required for the mother centriole anchoring to lipid membranes and notably the apical pole (Tanos et al., 2013). This suggested that the defective positioning we observed might result from a lack of anchor rather than a defective migration toward the apical pole. To distinguish these two possibilities we tracked centrosome 3D migration in centrin1-GFP expressing cells treated with either siRNA against Cep164 or non-targeting control siRNA. In Cep164 knocked-down cells, centrosome migration was severely defective. Few centrosomes did not take off at all from the basal pole, and the others could not reach the apical pole (Figure 2D), showing that the migration process itself, and not only the following anchoring step, was compromised. These results indicated that some primary ciliogenesis effectors participate in centrosome migration, notably Cep164, while others, such as IFT88, do not.

### Remodeling of the microtubule network during centrosome migration

We then investigated whether microtubules were involved in the regulation of centrosome migration upon starvation. The addition of nocodazole to depolymerize microtubules immediately after serum starvation blocked centrosome migration and ciliogenesis (Figure S3). When added five hours after starvation, ie after the completion of centrosome migration, it had no detectable effect on centrosome position and cilia elongation (Figure S3). This confirmed the specific implication of microtubules in centrosome migration. To investigate the role of microtubules in centrosome migration in more details, live imaging of micropatterned RPE1 EGFP-centrin 1 cells transduced with MAP4-RFP (Ganguly et al., 2013) to label the microtubules was carried out (Figure 3A and B, movie S1 and S2). Imaging of cells cultured in the absence of serum revealed that centrosome migration typically occurred 2 to 4 hours after serum starvation and this coincided with a dramatic increase in the number of microtubules surrounding the centrosome (Figure 3A). In cases where centrosome migration did not occur, no increase in microtubule density surrounding the centrosome was observed (Figure 3B). MAP4, being a known microtubule bundler, could be responsible for this effect. We controlled this by measuring the changes in microtubule density in non-transfected cells by fixing and staining serum-starved, micropatterned RPE1 cells with an antibody to α-tubulin (Figures 3C and D). Plotting of these measurements against the axial position of the centrosome showed that microtubule density increased significantly after serum starvation and there was a positive correlation between centrosome position and microtubule density (correlation coefficient of 0.9, Figure 3D). The monitoring of microtubule network reorganization during centrosome 3D migration showed that microtubule network densification was associated to the clustering of microtubule in a large bundle (movie S1). The orientation of this bundle between the centrosome and the basal pole suggested that it was exerting pushing rather than pulling forces. Indeed pulling forces would rather be associated to a bundle connecting the centrosome to the apical pole, as in the case of centrosome motion toward immune synapse (Yi et al., 2013). To quantify the occurrence of such pushing bundles, we fixed cells during the centrosome migration process, ie 80 minutes after serum starvation, and measured the frequency of bundles orientation toward basal or apical pole, whenever such a bundle could be detected (Figure 3E). Most centrosomes were associated to a microtubule bundle pointing toward the basal pole suggesting that pushing forces had a major contribution to centrosome propulsion toward apical pole.

**Figure 3.**
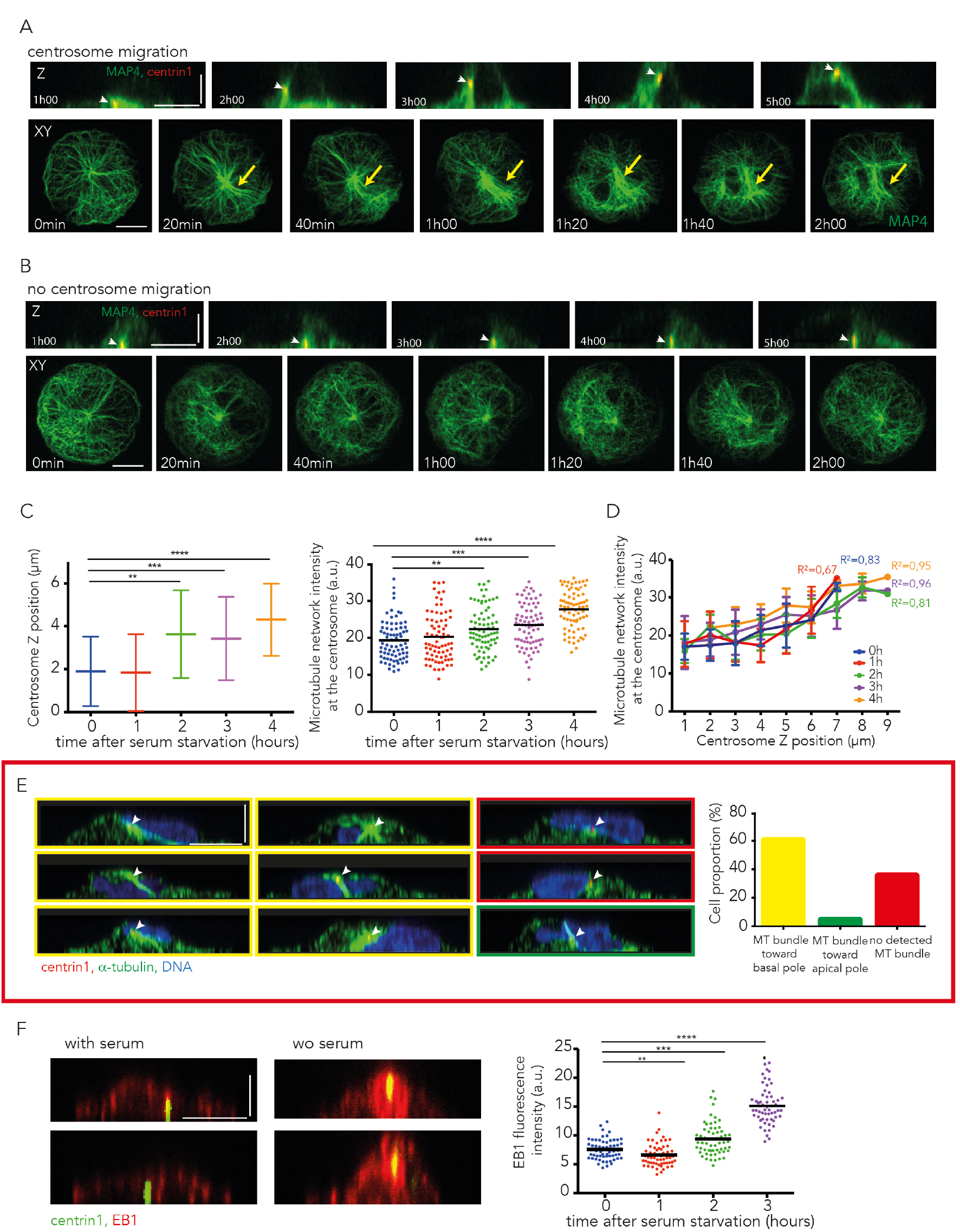
Microtubule network remodeling during centrosome migration. **A, B)** Microtubule network organizations were studied by time lapse imaging of RPE1 cells expressing EGFP-centrin1 (red) and MAP4-RFP (green). Two examples are shown, one where centrosome (indicated by white arrow heads) migrated to the apical pole (A) and another where it did not (B). Orthogonal and top views are shown. Microtubule network symmetry break and densification is shown with a yellow arrow. **C)** Measurement of centrosome Z position (left) and α-tubulin fluorescent intensity in a 5 µm box surrounding the centrosome (right) in thymidine-synchronized serum-starved RPE1 cells expressing EGFP-centrin1 for various delays after serum removal (n=75 cells per condition). **D)** The graph shows the microtubule network density at the centrosome against centrosome Z position at various time points after serum starvation in thymidine-synchronized RPE1 cells expressing EGFP-centrin1. In all conditions, the two parameters were correlated. **E)** RPE1 cells expressing EGFP-centrin1 (red, indicated by white arrows) were fixed 120 min after serum withdrawal and stained for α-tubulin (green) and DAPI (blue). The side views facilitated the visualization of microtubule bundles orientation and quantification of cell proportion exhibiting either a microtubule bundle toward basal pole (images and bar graph with yellow outlines) or a microtubule bundle toward apical pole (images and bar graph with green outlines) or no detected microtubule bundle (images and bar graph with red outlines) (results of 4 independent experiments, n=116 cells). **F)** Staining of serum-starved RPE1 cells expressing EGFP-centrin1 (green) with an antibody to EB1 (red). The graph shows EB1 fluorescence intensity measurements in a 5 µm box surrounding the centrosome (1 experiment, n=60 cells per condition). XY scale bars represent 10 µm and Z scale bars represent 5 µm.

Combined, these data suggested that increased microtubule nucleation and/or stabilization was responsible for a densification of the microtubule network that further generated the pushing forces required to move the centrosome to the apical surface. To confirm such a model, serum-starved RPE1 cells were stained with an antibody to the microtubule end-binding protein EB1 and its fluorescent intensity was measured over time (Figure 3F). Strikingly, EB1 levels were found to be nearly 2-fold higher at the centrosome after 3 hours of serum starvation, indicating that increased microtubule nucleation was likely responsible for driving centrosome migration during primary cilium formation.

### Microtubule stabilization drives centrosome movement

Microtubule stabilization could also participate in network densification and centrosome migration. Interestingly, serum starvation has been shown to reduce microtubule dynamics (Danowski, 1998) and increase characteristic post-translational modifications of stabilized microtubules during primary ciliogenesis (Berbari et al., 2013). To investigate whether such changes could specifically contribute to centrosome migration, we subjected micropatterned RPE1 cells (serum-starved for 1, 2 or 3 hours) to a brief cold-shock for 12 minutes and stained them with an antibody to α-tubulin (Figure 4A). Quantification of the polymerized tubulin amount present in the cell showed that there was a remarkable 2-fold increase in the number of cold-stable microtubules 2 hours after serum starvation. This increase in microtubule stability appeared to be transient, with tubulin levels decreasing after 3 hours of serum starvation but remaining significantly higher than those of serum-fed cells. These data support a model whereby increased microtubule nucleation and microtubule stabilization work synergistically to generate a dense network of stable microtubules upon which the centrosome can migrate.

**Figure 4.**
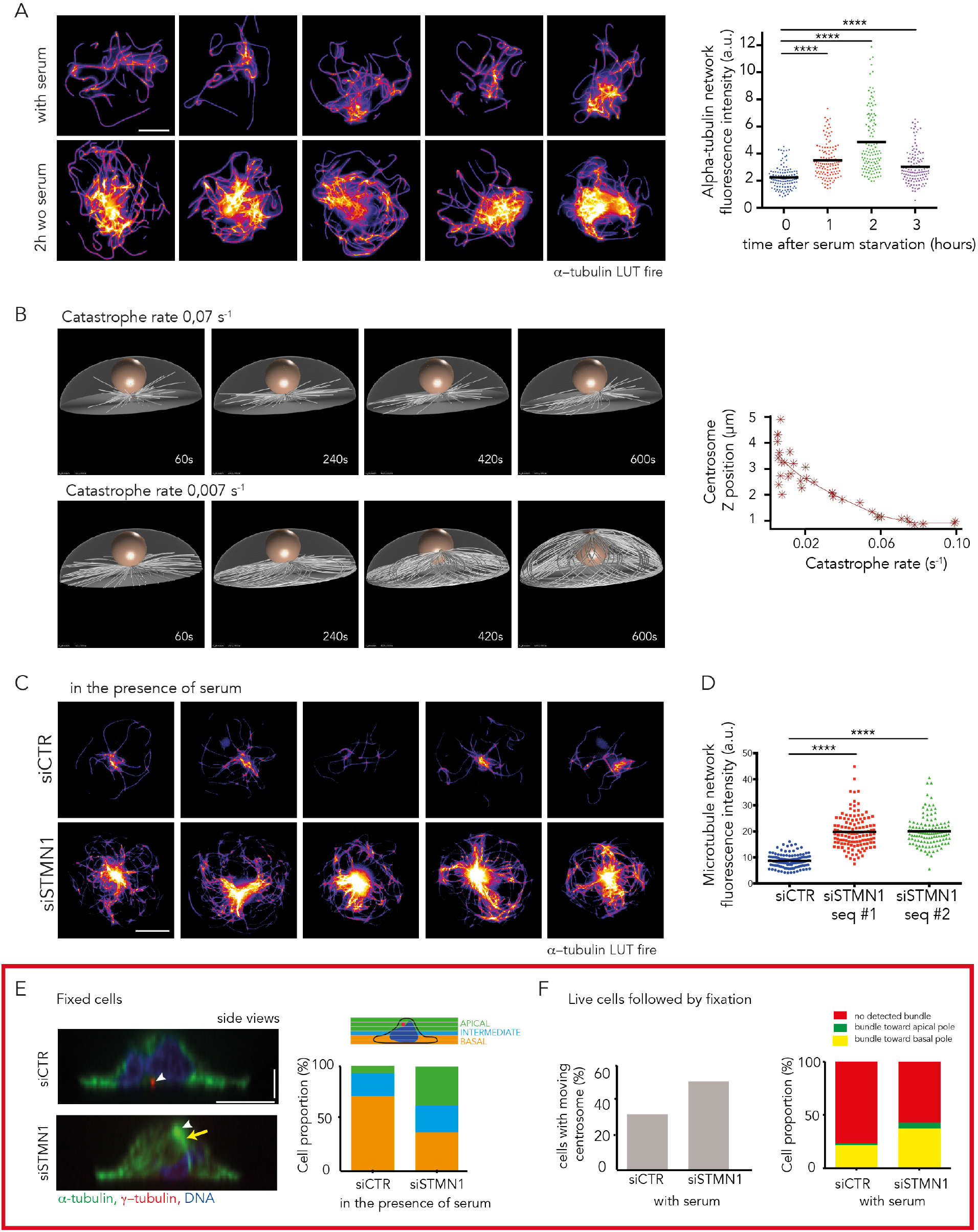
Microtubules stabilization after serum starvation promotes centrosome migration. **A)** Identification of cold-resistant microtubules. Serum-starved RPE1 cells were subjected to cold-shock, (on ice for 12 minutes) fixed and stained with an antibody to α-tubulin (fire LUT). Images show five examples of serum-starved and five examples of serum-fed cells (LUT fire). The graph shows measurements of α-tubulin fluorescence intensity after cold-shock for various delays after serum removal (results of 3 independent experiments, n=125 cells per condition). **B)** 3D numerical simulations of microtubule growth from the centrosome at the basal pole. They showed that longer microtubules, induced by reducing the catastrophe rate, induce a symmetry break in the network architecture that is capable of pushing centrosome to the apical surface. The graph shows the centrosome Z position according to catastrophe rate. **C, D)** Identification of cold-resistant microtubules in serum-fed cells treated with either control siRNA or siRNA against the tubulin sequestering protein stathmin 1. The same conditions as in (A). Images show five examples of serum-fed cells treated with control siRNA and five examples of serum-fed cells treated with stathmin 1 siRNA (C). The graph shows measurements of α-tubulin fluorescence intensity after cold-shock (results of 2 independent experiments, control siRNA, n=125 cells; stathmin 1 siRNA: 2 siRNA sequences, n=125 cells each) (D). **E)** Stathmin 1 was depleted by siRNA from RPE1 cells cultured in the presence of serum to promote microtubule growth and observe its effect upon centrosome position. Cells were fixed and stained for α-tubulin (green), γ-tubulin (red) and DNA (blue). Centrosomes are indicated by white arrows and microtubule network symmetry break and densification is shown with a yellow arrow (left side views images). The graph shows cell percentage displaying basal centrosome (located from 0 to 2 μm above the glass substrate), intermediate centrosome (between 2 and 3 μm above the glass substrate) and apical centrosome (located more than 3 μm above the glass substrate) (results of 3 independent experiments, control siRNA, n=100 cells; stathmin 1 siRNA: 2 siRNA sequences, n=150 cells each). **F)** Time lapse imaging for 80 min of serum-fed RPE1 cells expressing EGFP-centrin1 treated with control siRNA or siRNA against stathmin 1 then fixation / immunostaining for α-tubulin. The left graph shows the percentage of cells exhibiting moving centrosome toward apical pole in each condition (siCTR vs siStathmin 1). The right graph shows microtubule bundle orientation frequency for each condition (1 experiment, control siRNA: n=63 cells; siRNA against stathmin 1: n=45 cells). XY scale bars represent 10 µm and Z scale bars represent 2.5 µm.

We tested this model using 3D numerical simulations (Foethke et al., 2009) to investigate how microtubule network density could impact upon the axial position of the centrosome (Figure 4B). We built upon our previous simulations showing centrosome decentering upon microtubule lengthening in 2D (Letort et al., 2016; Burute et al., 2017) by taking into account cell shape in 3D and the centrosome's interaction with the nucleus. Catastrophe rate variations were used to modulate microtubule length and network density. Increasing microtubule length by reducing their catastrophe rate from 0.06 to 0.02 event per sec [range estimated from (Janson et al., 2003)] resulted in the formation of dense network of stable microtubules capable of generating sufficient force to push the centrosome to the apical surface (Figure 4B, movies S3 and S4). No minus-end directed motors capable of exerting pulling forces from the cell cortex were added to these simulations. These results indicated that a reduction in the catastrophe rate that resulted in the formation of a network of stable microtubules, as observed experimentally in serum-starved cells, could reorganize microtubule network architecture and the net orientation of pushing forces, so as to destabilize centrosome basal position and push it toward the apical pole.

To gather evidence to support the numerical simulation data, the tubulin-sequestering protein stathmin 1 (Belmont and Mitchison, 1996) was depleted from cells to increase the pool of free tubulin available and promote microtubule polymerization (Figures 4C and Supplementary S1, S2D). Imaging of the microtubule network in RPE1 cells treated with siRNA against stathmin 1 in presence of serum showed indeed a network of cold-stable microtubules reminiscent to those observed in serum-starved cells. The quantification revealed that tubulin levels were 2-fold higher after stathmin 1 depletion compared with siRNA control (Figure 4D). SiRNA-treated RPE1 cells were cultured in the presence of serum, fixed and stained with γ-tubulin and α-tubulin antibodies (Figure 4E). Confocal imaging and measurement of the axial position of the centrosome showed that it was significantly closer to the apical surface in the stathmin 1-depleted cells than the control siRNA-treated cells although serum had not been withdrawn in these experiments (Figure 4E). Furthermore, live imaging of centrin1-GFP labeled centrosomes followed by fixation and immuno-labelling of microtubules revealed an increase proportion of centrosomes moving toward the apical pole and an increased frequency of microtubule bundle pointing toward the basal pole in stathmin 1-knocked down cells compared to control cells in the presence of serum (Figure 4F). These results support the proposal that increased microtubule polymerization and stabilization is sufficient to generate a microtubule network capable of pushing centrosome toward cell apical pole.

### Actin network contraction and symmetry breaking promote apical centrosome motion

Numerous studies have implicated actin remodeling as part of the process of primary cilium formation (Kim et al., 2010; Pitaval et al., 2010; Dawe et al., 2009). We sought to characterize changes in actin cytoskeleton during primary ciliogenesis by staining RPE1 cells with phalloidin to label filamentous actin and α-tubulin antibody to label the microtubules (Figure 5A). Prior to serum starvation, the actin cytoskeleton of micropatterned RPE1 cells was observed to be radially symmetrical in agreement with previously published data (Tee et al., 2015). However, after 4 hours of serum starvation, the symmetry was broken and actin filaments appeared preferentially clustered to one side of the cell. The transverse arcs forming a ring of bundled filaments tend to contract toward an off-centered position (Figure 5A). This impacted upon the microtubule cytoskeleton, resulting in the asymmetric co-partitioning of microtubules with F-actin. This reorganization compressed the nucleus, which became higher and less spread. The symmetry break in the actin network forced the nucleus to be displaced from the center (Figure 5B).

**Figure 5.**
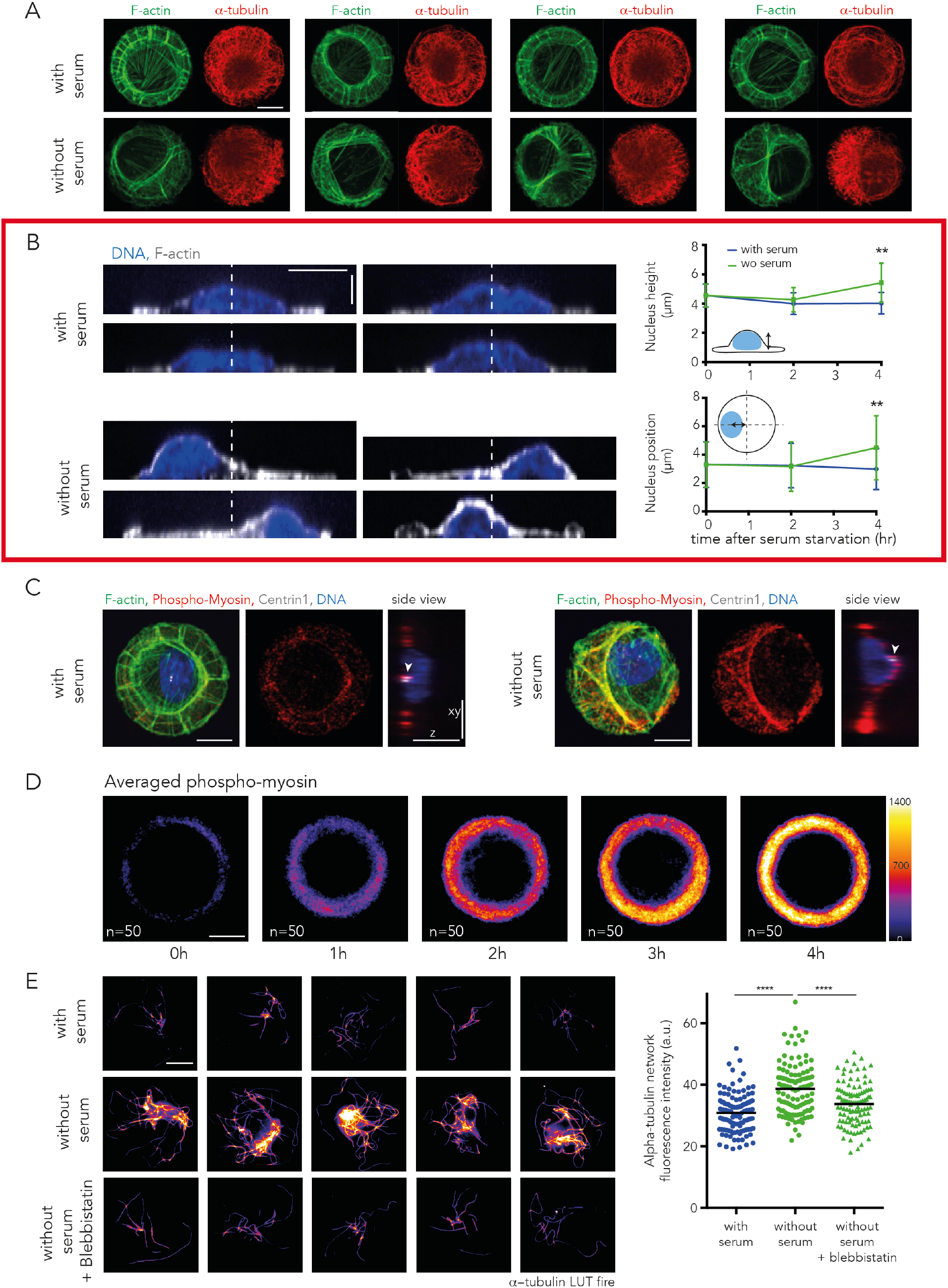
Contractility increase breaks actin cytoskeleton symmetry and promotes microtubule stabilization. **A)** Reorganization of the actin and microtubule cytoskeletons upon serum starvation. RPE1 cells were fixed 4h hours post serum withdrawal, stained with phalloidin to visualize F-actin (green) and immuno-stained with antibodies against α-tubulin (red) and compared to serum-fed cells. Images show four examples of serum-starved and serum-fed cells. **B)** Nucleus positioning upon serum starvation. RPE1 cells were fixed 4h post serum starvation, stained with phalloidin (gray) and DAPI (blue) and compared to serum-fed cells. Images show four examples of serum-starved and serum-fed cells. Gray dotted lines indicate cell symmetry axis. The upper graph shows measurements of nucleus height, the lower graph shows measurements of nucleus position in XY plane in fed and starved-cells at various time points after serum starvation (results of 2 experiments, n=75 cells). **C)** Immuno-staining against phospho-myosin (red) showed an intense staining along non-circular actin bundle (green) in serum-starved cells. White arrows point at centrosomes detected with anti-centrin1 antibodies (white). RPE1 cells were fixed 4h hours post serum withdrawal **D)** Averaging of phospho-myosin fluorescent intensity levels (fire LUT), obtained by stacking and averaging 50 images per condition, showed that the myosin phosphorylation increased after serum starvation. **E)** Identification of cold-resistant microtubules in serum-fed and serum-starved cells in the presence or absence of the myosin-II ATPase inhibitor blebbistatin. RPE1 cells were fixed 2h hours post serum withdrawal. Images show five examples of serum-starved, five examples of serum-fed cells and five examples of serum-starved cells treated with blebbistatin for 2h (LUT fire). The graph shows measurements of α-tubulin fluorescence intensity after cold-shock (results of 2 independent experiments, n=110 cells; for each condition). XY scale bars represent 10 µm and Z scale bars represent 5 µm.

To ascertain if myosin II was involved in actin remodeling during primary ciliogenesis, RPE1 cells were stained with phalloidin and phospho-MLC II antibody (Figure 5C). After serum starvation, actin filaments were found to be decorated with phospho-MLC II antibody, suggesting that remodeling of the actin cytoskeleton was due to myosin II activity. Staining of RPE1 cells with phospho-myosin antibody followed averaging of the fluorescent signal indicated that the level of phosphorylated myosin II increased with time after the induction of primary ciliogenesis (Figure 5D). While these data suggested that myosin II activity was required for actin remodeling, they did not provide direct evidence of a role for myosin II-dependent contractility in centrosome migration. To test for such a role, RPE1 cells were treated with the myosin II inhibitor blebbistatin and MT cold-resistance assays carried out. A dramatic reduction in the number of cold-stable microtubules, compared with controls, was observed after blebbistatin treatment, and indicated that myosin II-dependent contractility was involved in microtubule reorganization and stabilization (Figure 5E).

### Apical pole maturation follows centrosome migration

These results suggested the implication of an internal symmetry break in cytoskeleton organization that contrasted with the more classical view of centrosome being off-centered by the action of localized pulling forces from a defined portion of the cell cortex (Tang and Marshall, 2012; Barker et al., 2015). We challenged our interpretation by looking at classical markers of apical pole that could be involved in the local activation of pulling forces (ERM : ezrin, radixin, moesin, NuMA and p150Glued). Increased phosphorylation of ERM was observed after serum starvation (Supplementary Figure S4A). However, this increase could only be detected after the centrosome migration process, suggesting that it was a more downstream event. Increased recruitment of the dynein-interacting proteins p150Glued and NuMA to the apical cortex was also observed after serum starvation but was initiated only four hours after serum withdrawal when most centrosomes had already reached their apical position (Supplementary Figures S4B,C), further confirming that their local accumulation at the apical pole followed rather than promoted centrosome migration.

### Investigation of the role of ciliogenesis effectors in centrosome migration

The above data suggested that centrosome movement was driven by concomitant, and likely related (Joo and Yamada, 2014; Rao et al., 2014) stabilization of microtubules and contraction of the actin network, allowing symmetry breaks in both networks and the efficient production of microtubule-based pushing forces on the basal pole. Within this context, we decided to revisit the effect of the depletion of ciliogenesis effectors and test whether they affected centrosome migration via the mechanism we hypothesized. To that end, we investigated in more detail the cytoskeleton organizations resulting from the depletion of Cep164 and IFT88, because their ablation had opposing effects upon centrosome migration. In a microtubule cold-resistance assay, Cep164-depleted cells had fewer cold-resistant microtubules upon serum starvation (Figure 6A) whereas control and IFT88-depleted cells had more (Figures 6A,B and Supplementary Figure S5A). This suggested that the centrosome migration defect previously observed in Cep164-depleted cells was actually due to a failure to stabilize microtubules and remodel the network architecture accordingly. The proper centrosome migration in IFT88-depleted cells was consistent with proper microtubule reorganization in those cells. In parallel, phospho-myosin density appeared reduced in Cep164-depleted cells compared with control or IFT88-depleted cells (Figure 6C), further confirming the specific implication of the mechanism we discovered in the control of centrosome migration. Finally, the level of NuMA at the apical pole of serum starved cells, which was seen to rise few hours after centrosome migration, was lower in Cep164- and higher in IFT88-depleted cells compared to controls (Figure S5B). As Cep164 is a centrosome regulator with no described effect in cortical actin, the absence of accumulation of NuMA at the apical pole is likely to be a consequence of the defective centrosome migration. Altogether, these results supported our conclusion that apical pole maturation in these conditions was a consequence rather than the cause of centrosome migration.

**Figure 6.**
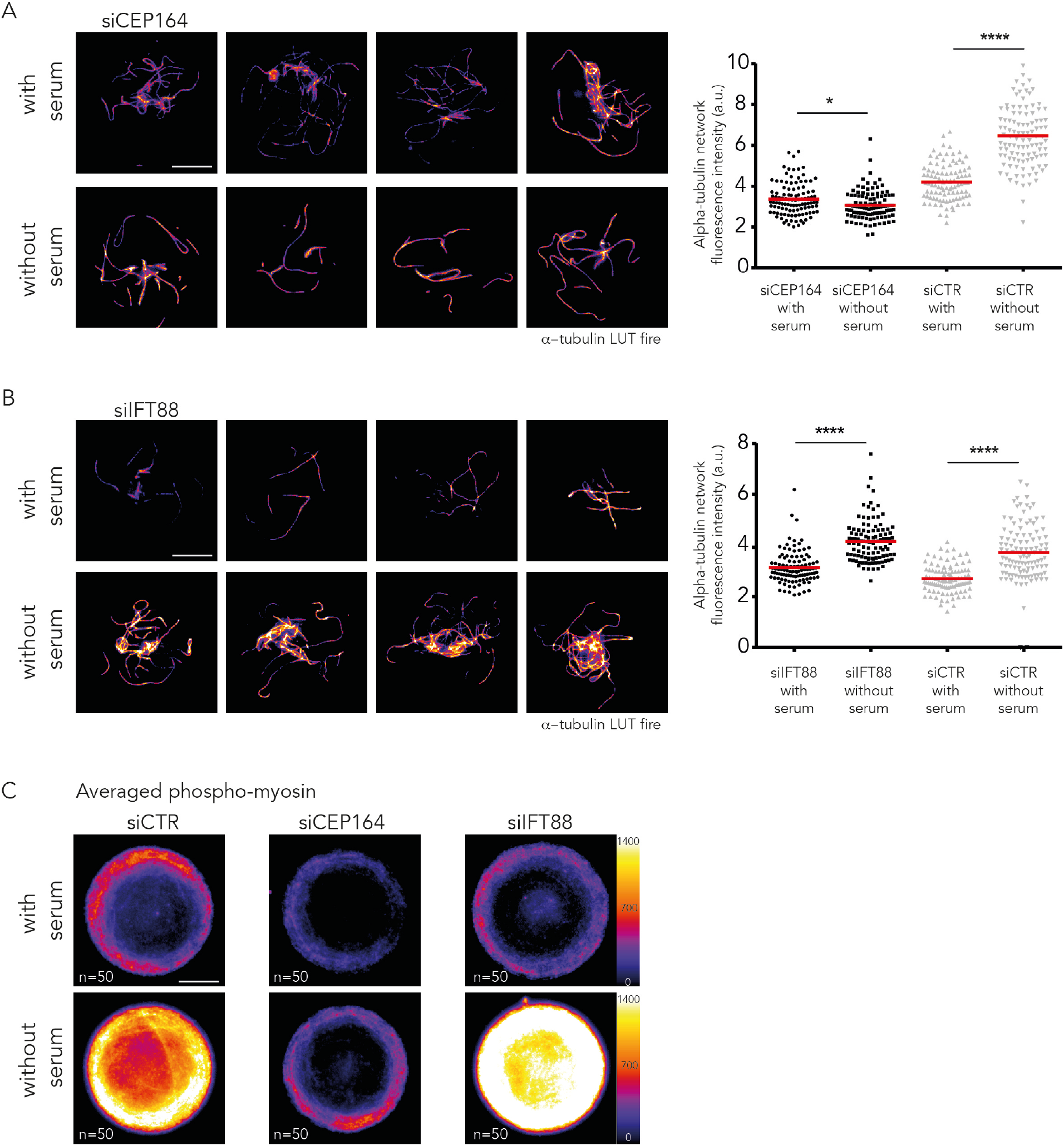
The ciliogenesis effector Cep164 affects microtubule stabilization and acto-myosin contractility upon serum starvation. **A)** Identification of cold-resistant microtubules in serum-starved and serum-fed cells treated with siRNA against Cep164. Images show four representative examples of Cep164-depleted serum-starved and Cep164-depleted serum-fed cells (experiments obtained with two distinct Cep164 siRNA sequences). Cells fixed after a brief cold-shock of 12 minutes following 3h of serum starvation. The graph shows measurements of α-tubulin fluorescence intensity after cold-shock (LUT fire) (results of 2 independent experiments, n=110 cells per condition). **B)** Same as in A with siRNA against Ift88 (results of 2 independent experiments, n=110 cells per condition). **C)** Averaging of phospho-myosin fluorescent intensity levels (fire LUT), obtained by stacking and averaging 50 images per condition. Cells were fixed four hours after serum starvation. Averaged images showed that myosin phosphorylation increased in control and Ift88-depleted cells, but not in Cep164-depleted cells. XY scale bars represent 10 µm.

## Discussion

In this paper we have exploited the technique of cell micropatterning to characterize in detail the previously poorly defined step of centrosome migration during primary ciliogenesis. Microtubules in cells undergoing primary ciliogenesis were found to be more resistant to cold treatment, suggesting that they were more stable. In addition, increased levels of EB1 and tubulin at the centrosome were also observed. Elevated EB1 levels at the centrosome could represent an increase in microtubule nucleation or anchoring, as EB1 is involved in both (Yan et al., 2006). Numerical simulations in 3D showed an interesting consequence of this remodeling of the microtubule network. Increasing microtubule stability appeared sufficient to force network reorganization, leading to a vortex-like conformation that destabilizes the basal position of the centrosome and pushes it up toward the cell apical pole. This mechanism was further confirmed experimentally, using the depletion of the tubulin sequestering protein stathmin 1 to increase the level free tubulin available for incorporation into polymer to generate an array of long microtubules. This resulted in the formation of an array capable of transmitting sufficient force to push the centrosome to the apical surface in the absence of any of the other compounding effects associated with serum starvation. Furthermore, ciliogenesis effectors involved in the regulation of centrosome migration, such as Cep164, also impacted microtubule stabilization and cell contractility, further supporting the implication of cytoskeleton remodeling in centrosome migration. This mechanism, relying on a symmetry break in the spatial organization of pushing forces, contrasts with the previously described mechanisms of centrosome off-centering in which unbalanced forces result from the asymmetric distribution cortical pulling forces (Morin and Bellaïche, 2011; Tang and Marshall, 2012).

These observations led us to propose the following speculative scenario for the induction of centrosome migration when cells enter quiescence (Figure 7). Centrosome maturation is initiated in the first two hours following serum starvation and primes ciliogenesis (Westlake et al., 2011; Lu et al., 2015). Although we have no evidence that this maturation directly impacts microtubule dynamics, the lack of microtubule stabilization in response to Cep164 knock-down suggests that the Cep164-dependent recruitment of the ciliary vesicle and associated components (Cajánek and Nigg, 2014; Schmidt et al., 2012) contributes to microtubule nucleation and stabilization. Microtubule lengthening appeared sufficient to induce a symmetry break in the spatial arrangement of microtubules. In parallel, microtubule stabilization is likely to feedback to acto-myosin contractility via specific kinases and phosphatases co-regulating the two pathways such as myosin phosphatase (Joo and Yamada, 2014; Rao et al., 2014). Increase in acto-myosin activity is sufficient to break the symmetry of the contracting network as previously observed in several contractile systems (Sedzinski et al., 2011; Paluch et al., 2005; Yam and Theriot, 2004). This asymmetric actin flow can move the nucleus away from the cell center (Gomes et al., 2005) and thereby facilitates centrosome rise from its initial central position and further contributes to the asymmetric reconfiguration of the microtubule network that was initiated by microtubule stabilization. Furthermore, acto-myosin contractility further contributes to microtubule stabilization. The increase in pushing forces associated to microtubule polymerization (Laan et al., 2008) and the imbalance in microtubule distribution destabilize the centrosome's position at the basal pole (Pinot et al., 2009; Letort et al., 2016) and push it toward the apical pole, allowing microtubule elongation and the release of elastic stress accumulated as a result of their bent conformation. When the centrosome reaches the apical membrane, it brings along minus-end directed motors, such as dyneins, and their associated proteins like NuMA (Merdes et al., 1996), which can then interact with the plasma membrane (Kotak et al., 2014). This local accumulation of microtubule interacting proteins further contributes to the anchoring of the centrosome at the apical pole and the consequential accumulation of centrosome-associated proteins as well as cargos transported along the microtubules, which contribute to later stages of ciliogenesis (Reiter et al., 2012). Although our observations do not exclude a possible contribution of pulling forces exerted on centrosomal microtubule by minus-end directed motors anchored to a nascent apical pole, they provide compelling evidence for a major role played by microtubule pushing on the basal pole.

**Figure 7.**
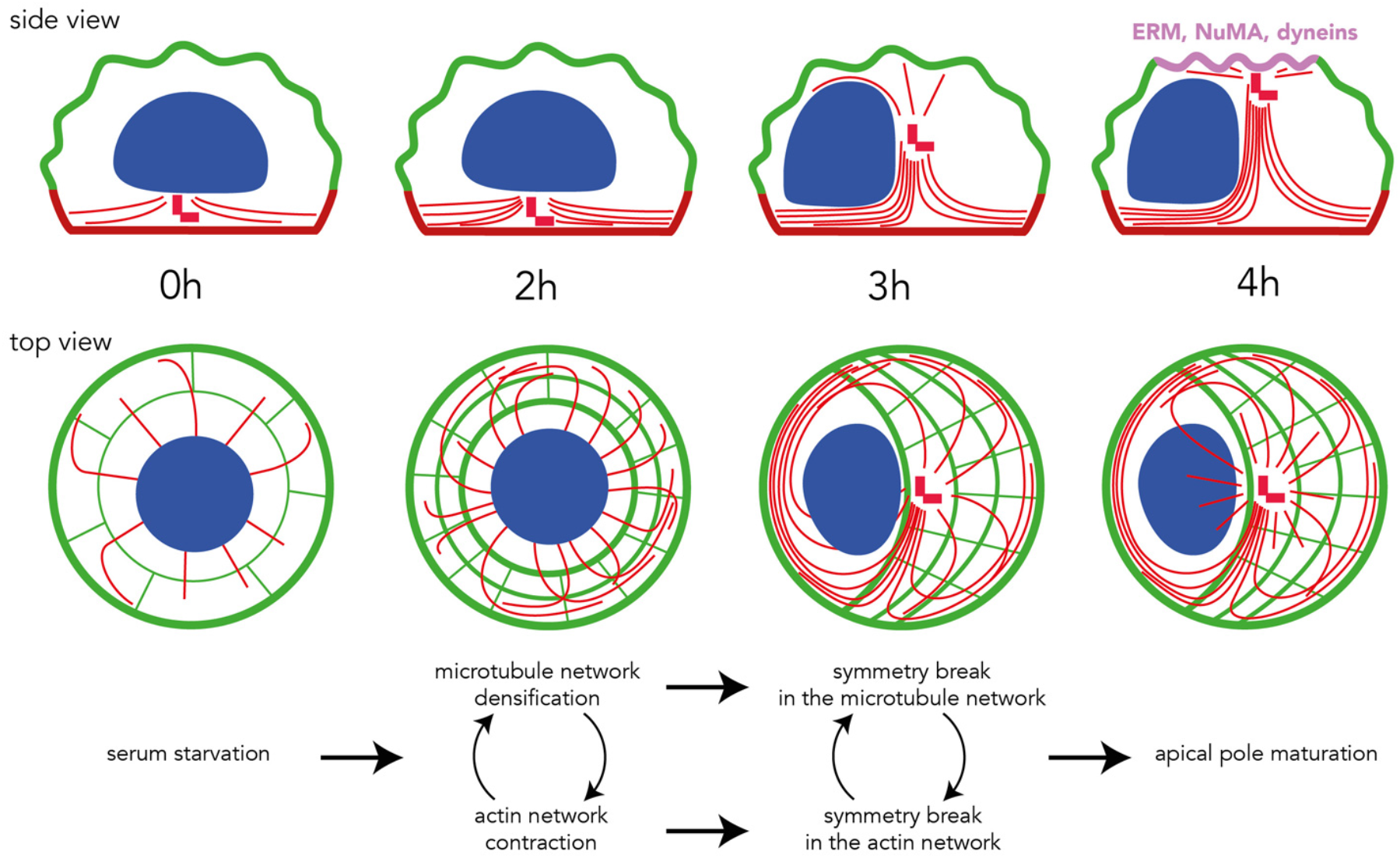
Proposed sequence of events driving centrosome migration to cell apical pole. These schemes show side and top view of cytoskeleton rearrangements following serum withdrawal. Microtubule network densification and actin network contraction break symmetry of both networks, which results in the production of pushing forces moving the centrosome to the dorsal surface. Upon contact, the centrosome promotes local surface maturation into an apical pole and centriole anchoring and elongation to form the primary cilium.

The key step in the mechanism we described is the transient stabilization of microtubules upon serum starvation. Serum starvation or activation of Rac1 or Cdc42 are known to result in microtubule end stabilization and lifespan increase (Grigoriev et al., 2006; Danowski, 1998) but the underlying mechanism still remains to be uncovered. Interestingly, microtubule stabilization and bundling upon entry into quiescence has been described in other systems. Recent work in fission yeast has shown that upon entry into quiescence S. pombe assembles its microtubules into a single bundle that is attached to the spindle pole body, the yeast equivalent of the centrosome (Laporte et al., 2015). The assembly of this single microtubule bundle from the three to five bundles of microtubules that are normally present during interphase is Ase I and Mto I dependent. Interestingly, homologs of these proteins exist in humans and they are PRC1 and CDK5RAP2 (Cep215), respectively. PRC1 is a microtubule bundling protein that binds to the microtubules of the central spindle that forms in late mitosis (Mollinari et al., 2002), while CDK5RAP2 is a pericentriolar material protein (PCM) involved in the nucleation of microtubules through the stimulation of γ-tubulin ring complex activity (Choi et al., 2010). It would be interesting to determine if CDK5RAP2 participates in the increased nucleation of microtubules during centrosome migration and if PRC1 is involved in reorganizing the microtubule network during primary cilium formation. The exact role of other centrosomal proteins, such as Cep164, and the associated recruitment of the ciliary vesicle in microtubule network reorganization during early ciliogenesis is still obscure. Interestingly epithelial to mesenchymal transition was recently shown to be associated with a reduction in microtubule number and stability that was required for cell migration (Burute et al., 2017). In parallel, the simple knockdown of Cep164 was shown to be sufficient to induce such a transition and to foster the migration of epithelial kidney cells (Slaats et al., 2014). These two observations further support a probable role for Cep164 in the regulation microtubule nucleation and stabilization that definitely deserves further investigation.

Our data also suggest a role for myosin-based contractility in reorganizing the actin cytoskeleton which seems to facilitate centrosome migration by moving the nucleus and force microtubule network asymmetry. Reorganization of the actin cytoskeleton from a radially symmetrical array to an asymmetrical one occurred within 3 hours of the induction of primary ciliogenesis and was abolished by treatment with the myosin II inhibitor blebbistatin (data not shown). Myosin II and ROCK inhibition were shown previously to impair centrosome migration (Pitaval et al., 2010). Here we confirmed earlier observations that the knock-down of meckelin, emerin and nesprin2, which are actin-binding proteins ensuring the centrosome-nucleus connection, perturbs centrosome migration (Dawe et al., 2009). Interestingly this suggests that the centrosome-nucleus link needs to be maintained during migration. Nucleus deformation or rotation upon acto-myosin contraction may help apical centrosome displacement. The nucleus could also act as a guide to orient the pushing forces produced by the microtubule network. Our observations add to the increasing knowledge about the implication of acto-myosin contractility in ciliogenesis but the exact mechanism remains to be established.

Altogether, our data demonstrate that centrosome migration to the apical surface is orchestrated by coordinated changes in the actin and microtubule cytoskeletons with an increase in microtubule stability playing an important part in the process. Identifying the factors responsible for mediating microtubule stability in response to serum starvation and the connection with the actin cytoskeleton remodeling should allow us to further understand this major intracellular reorganization occurring when cells enter quiescence.

## Materials and Methods

### Cell Culture

Human telomerase-immortalized retinal pigment epithelial 1 (RPE1) cells (Clontech) and RPE1 cells stably expressing EGFP-centrin1 (a kind gift of Alexey Khodjakov) or Lifeact-GFP were cultured in a humidified incubator at 37°C in DMEM/F12 medium supplemented with 10% fetal bovine serum and 1% penicillin/streptomycin (all from Life Technologies).

### Cell plating on Micropatterns slides

Disc-shaped micropatterned coverslips were obtained from CYTOO or produced in house according to previously established protocols (Azioune et al., 2009)

### Inhibitors

RPE1 cells were treated with 50 μM blebbistatin for 2 hours in the absence of serum. Synchronization of RPE1 cells was carried out using a double thymidine block, culturing the cells in medium containing 2 mM thymidine (Sigma) for 16 hours, releasing for 10 hours, and culturing again in thymidine-containing medium for a further 16 hours. Cells were released from the block by removing the thymidine-containing medium and after 10 hours, when the cells were in early G1, they were plated onto micropatterns.

### Viral transduction

RPE1 cells were transduced with BacMam MAP4-RFP virus (Life Technologies) according to the protocol provided.

### Small-interfering RNA treatment

RPE1 cells were transfected with siRNAs (Qiagen and Dharmacon) using Lipofectamine RNAi Max transfection reagent (Life Technologies) at a final concentration of 10 nM following the manufacturer's protocol. At least two independent siRNAs were tested for each target and two or three independent experiments.

### Antibodies and cytoskeletal labeling agents

Primary antibodies used in this study were obtained from the following sources and used at the following dilutions: mouse anti-acetylated tubulin (Sigma clone 6-11B-1; 1/10,000 for IF), rabbit anti-α−tubulin (Serotec AbD MCA77G; 1/3,000 for IF), rabbit anti-Cep164 (Erich Nigg; 1:2,000 for WB), mouse anti-EB1 (BD Biosciences #610535; 1/500 for IF), rabbit anti-γ-tubulin (Abcam ab11317; 1/1,000 for IF), rabbit anti-GAPDH (Santa Cruz #25778; 1/2000 for WB), rabbit anti-IFT88 (Proteintech #13967; 1/200 for WB), mouse anti-lamin A/C (Sigma, clone 4C11; 1/5,000 for WB), rabbit anti-NuMA (Santa Cruz #48773; 1/100 for IF), mouse anti-p150Glued (BD Biosciences #612709; 1/100 for IF), anti-pERM (Cell Signaling Technology #3141; 1/800 for IF), rabbit anti-stathmin 1 (Abcam #52630; 1/50,000 for WB) and rabbit anti-phospho myosin light chain 2 (Ser19) (Cell Signaling Technology #3671; 1/50 for IF). Alexa fluorophore-conjugated secondary antibodies (Molecular Probes) were diluted 1/1,000. Alexa fluorophore-conjugated phalloidin (Molecular Probes) was re-suspended in methanol and diluted 1/500 in PBS.

### Immunofluorescent staining

Different fixation protocols were used and depended on the antigen-binding characteristics of the antibody. For the siRNA screening experiments, where cells were stained with γ−tubulin and acetylated tubulin antibodies, fixation was carried out using cold methanol/acetone (50/50) on ice for 5 minutes. EB1 staining required the cells to be fixed with cold methanol for 5 minutes. Phosphorylated myosin light chain 2 antibody staining required pre-permeabilization of the cells with 0.1% Triton X-100 (Sigma) in MTBS buffer (60mM PIPES, 25 mM Hepes, 5mM EGTA, 1 mM MgCl, pH7) prior to fixation with 4% PFA for 15 minutes at ambient temperature. Stainings for phosphorylated ERM proteins, p150Glued and NuMA were carried out after fixation with 4% paraformaldehyde for 15 minutes, followed by permeabilization with 0.1% Triton X-100 in PBS for 3 minutes and incubation with the antibody overnight at 4°C for pERM. Where cells were stained with α-tubulin antibody and phalloidin, fixation was carried out with 0.5% glutaraldehyde in MTBS buffer (60mM PIPES, 25 mM Hepes, 5mM EGTA, 1 mM MgCl, pH7) containing 0.1% Triton X-100 for 15 minutes followed by quenching with NaBH4 for 10 minutes, both at ambient temperature.

For all conditions, after fixation, the cells were washed then blocked with PBSA (PBS containing 1.5% bovine serum albumin (Sigma) for 30 minutes. The cells were stained with primary antibodies diluted in PBSA for 1 hour, with the exception of the pERM where the cells were stained overnight at 4°C, followed by extensive washing with PBSA and staining with secondary antibodies diluted in PBSA for 30 minutes. The cells were washed with PBS and the DNA labeled with 0.2 μg/ml DAPI (Sigma) for 1 minute. After washing the cells with water, the coverslips were air-dried and mounted onto slides using Mowiol.

### Cold resistance assay

Cells were subjected to a cold shock on ice for 12 min and then pre-permeabilized with MTBS buffer (60mM PIPES, 25 mM Hepes, 5mM EGTA, 1 mM MgCl, pH7) containing 0.1% Triton X-100 for 20 seconds and fixed with 0.2 % glutaraldehyde in the same buffer for 15 min.

### Live and fixed cell acquisition and analysis

Live cell imaging was carried out on a Nikon spinning disk microscope equipped with a 60X, 1.4 NA objective lens and a HQ2 CoolSnap camera (images were taken every 20 min). Images of fixed cells were captured on an Olympus BX61 microscope equipped with a 100X, 1.4 NA objective lens and a HQ2 CoolSnap camera. Some fluorescent images shown are maximal projections of Z stacks acquired with oil immersion objectives at 100× (NA = 1.4) mounted on a piezo ceramic (Physics Instruments). Both microscopes were controlled with Metamorph software (MDS Analytical Technologies). Fluorescence images were also taken using confocal Z stacks acquired with a confocal microscope (TCS-SP2; Leica) through a 63× objective (NA = 1.4).

Automated image acquisition and analysis was performed as previously described (Pitaval et al., 2013). For the siRNA screen, centrosome Z position, percentage of ciliated cells and cilia length are quantified for each treated cell.

### Western Blotting

Proteins were separated by SDS polyacrylamide gel electrophoresis and transferred onto nitrocellulose membrane using a semi-dry Western blotting apparatus (BioRad). The membranes were blocked with PBS containing 5% non-fat milk for 1 hour at ambient temperature. After blocking the membranes were probed with primary antibodies overnight at 4°C. The membranes were washed four times with blocking buffer before adding HRP-conjugated secondary antibodies (Life Technologies), diluted as recommended in blocking buffer, and incubating for 30 minutes at ambient temperature. After washing three times with PBS containing 0.1% Tween-20 (Sigma) the membranes were developed using ECL reagent (Life Technologies) and imaged on the ChemiDoc system (BioRad) or by exposing to scientific imaging film (Kodak).

### Numerical simulations

Numerical simulations were carried out using Cytosim software (Nedelec and Foethke, 2007).

### Statistical tests

A Fisher exact test was performed to analyze contingency tables comparing the number of cells in two conditions for both control and siRNA, using R version 3.3.2. All the other data were presented as mean ± standard deviation (SD). Results were analyzed using Mann-Whitney test (GraphPad Prism). ****p<0.0001; ***p<0.001; **p<0.01; *p<0.05.

## Acknowledgements

We thank Alexey Khodjakov for the kind gift of RPE EGFP-centrin1 cells, Erich Nigg for anti-Cep164 antibody, Laurent Blanchoin and Maxence Nachury for helpful discussions. This work was supported by an ERC Starting grant to MT (SPiCy 310472).

## Supplementary Figures

**Figure S1.**
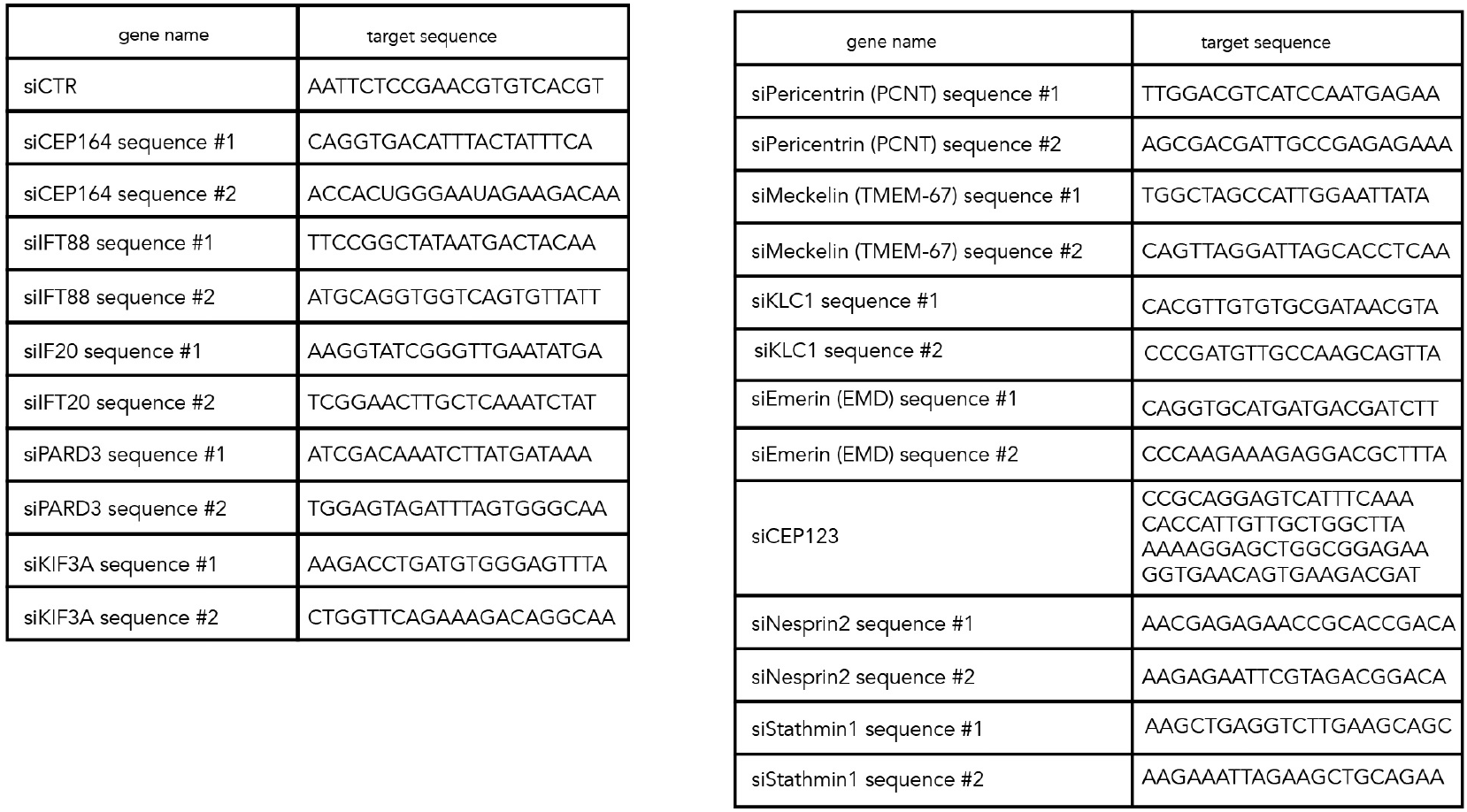
Sequences used in the study. The sequences used in the candidate-based siRNA screen.

**Figure S2.**
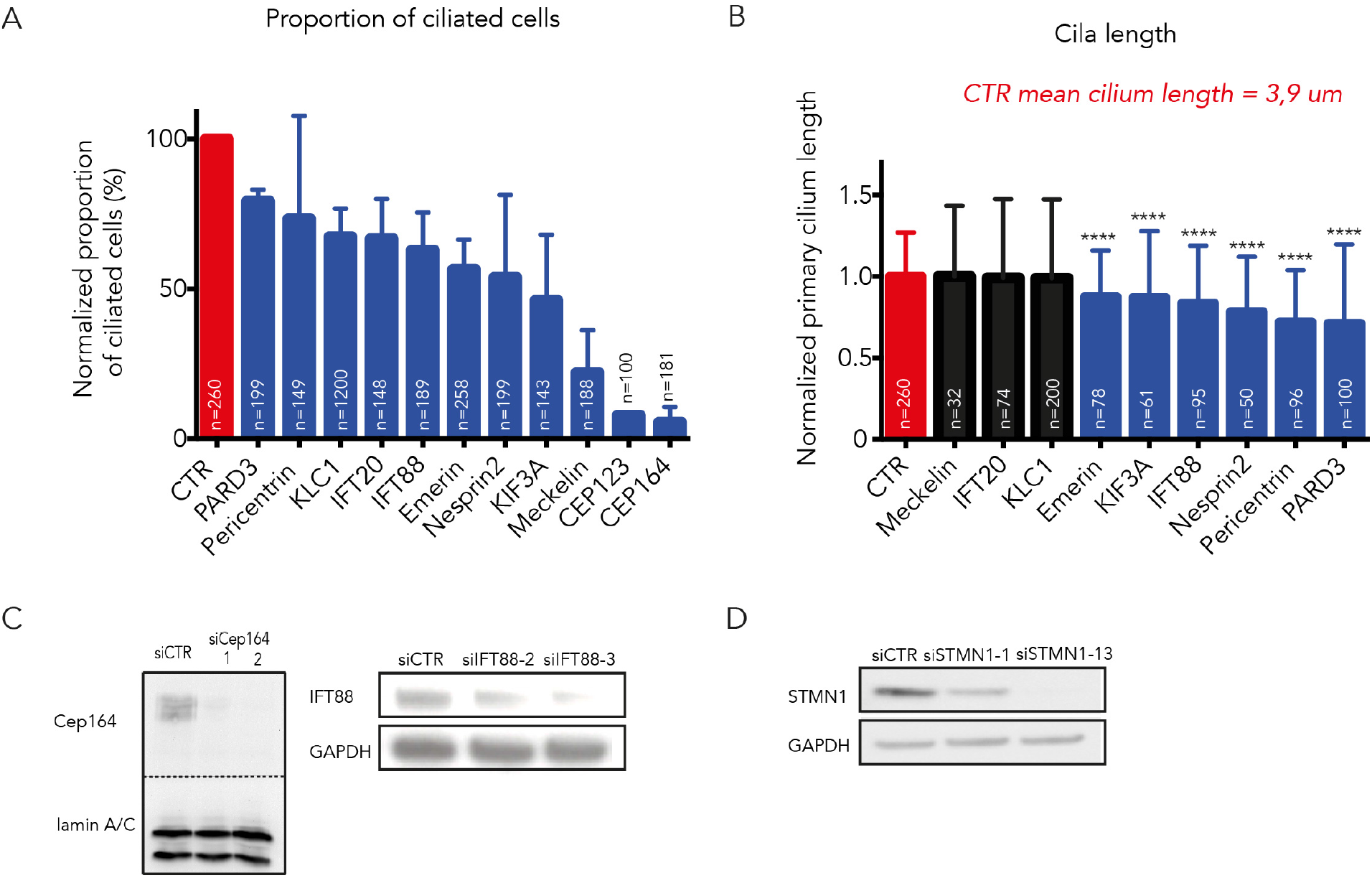
Validation of siRNAs used in the study. **A)** Control of siRNA efficiency presented in Figure 2A by quantification of the number of ciliated cells. Proportions were normalized with respect to non-targeting control siRNA for each condition. Red color is the control, blue means there is a statistically significant difference with respect to control. Values in the table represent the number of independent experiments used for the siRNA screen (n indicated at the bottom of each table), the percentage of ciliated cells for independent experiments and for the two siRNA sequences. **B)** Control of siRNA efficiency presented in Figure 2A by quantification of cilia length. Proportions were normalized with respect to non-targeting control siRNA for each condition. Red color is the control, blue means there is a statistically significant difference with respect to control, black means there is no significant difference with the control. **C)** Western blots demonstrating the efficacy of the Cep164, Ift88 siRNAs. Lamin A/C or GAPDH antibodies were used as loading controls for the blots. **D)** Western blots demonstrating the efficacy of the stathmin 1 siRNAs. GAPDH antibodies was used as loading control for the blots.

**Figure S3.**
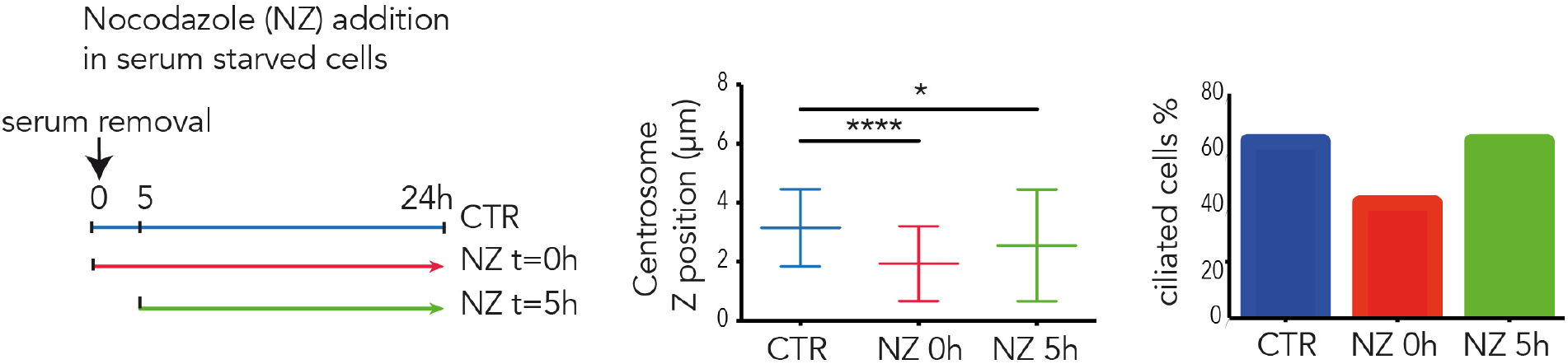
Nocodazole experiment. Effect of microtubule depolymerization on centrosome Z position and ciliogenesis rate by addition of nocodazole synchronously with (red) or 5 hours after (green) serum withdrawal (results of 2 independent experiments, n=75 cells per condition). Centrosome Z position (middle) and ciliated cells percentage (right) are represented for each condition.

**Figure S4.**
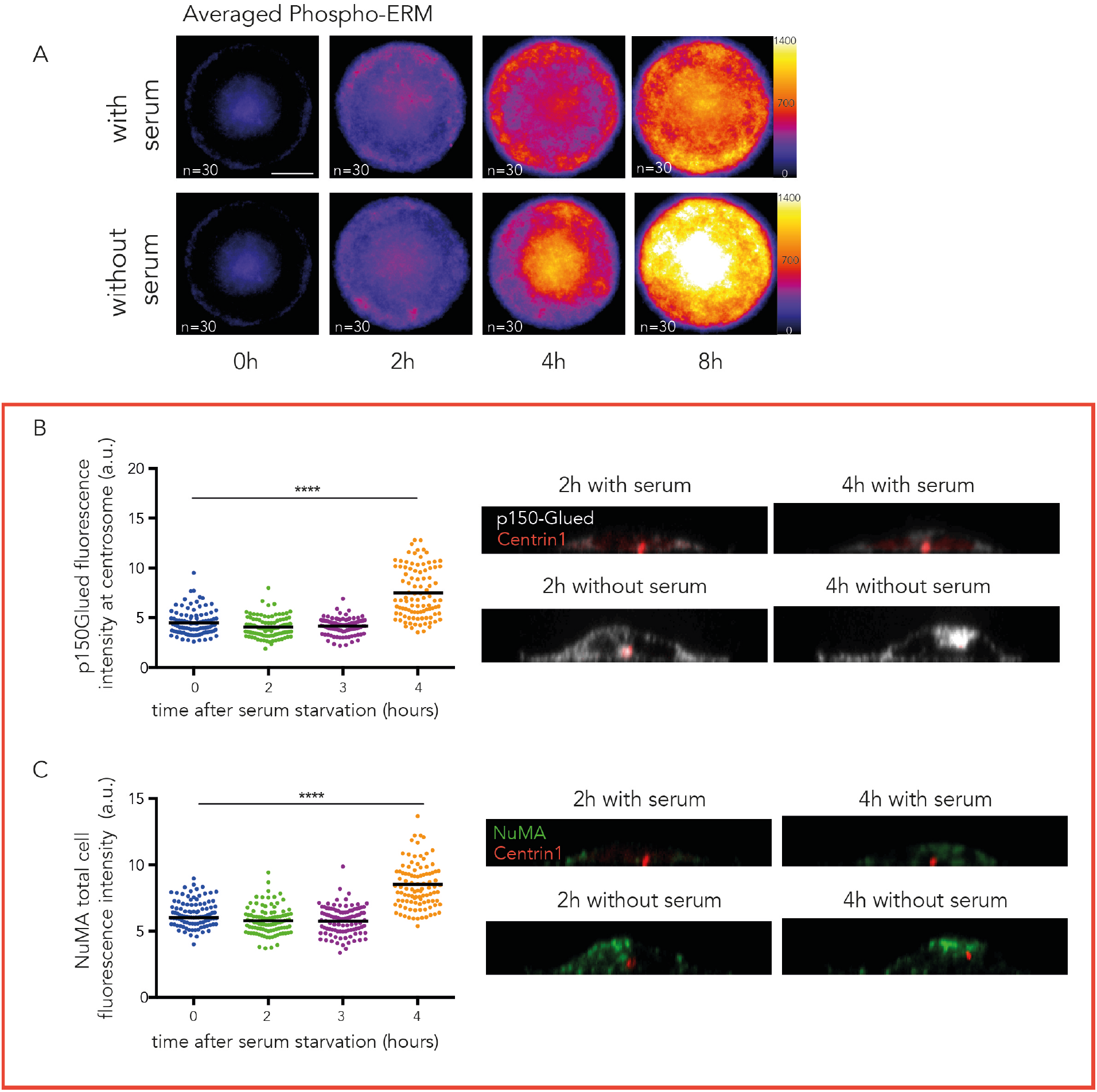
Recruitment of apical markers during centrosome migration. **A)** Averaging of phospho-ERM fluorescent intensity levels (LUT fire) in serum-fed and serum-starved cells obtained by stacking and averaging 30 images per condition. **B)** The graph shows the measurements of p150Glued fluorescence intensity at the centrosome in RPE1 EGFP-centrin1 cells following serum starvation (results of 2 independent experiments, n=100 cells per condition). Side views of RPE1 cells expressing EGFP-centrin1 (green), cultured in the presence or absence of serum, stained with antibody to p150Glued (grey). **C)** The graph shows the measurements of NuMA fluorescence intensity at the centrosome in RPE1 EGFP-centrin1 cells following serum starvation (results of 2 independent experiments, n=100 cells per condition). Side views of RPE1 cells expressing EGFP-centrin1 (green), cultured in the presence or absence of serum, stained with antibody to NuMA (red) (same cells than B). XY scale bars represent 10 µm and Z scale bars represent 5 µm.

**Figure S5.**
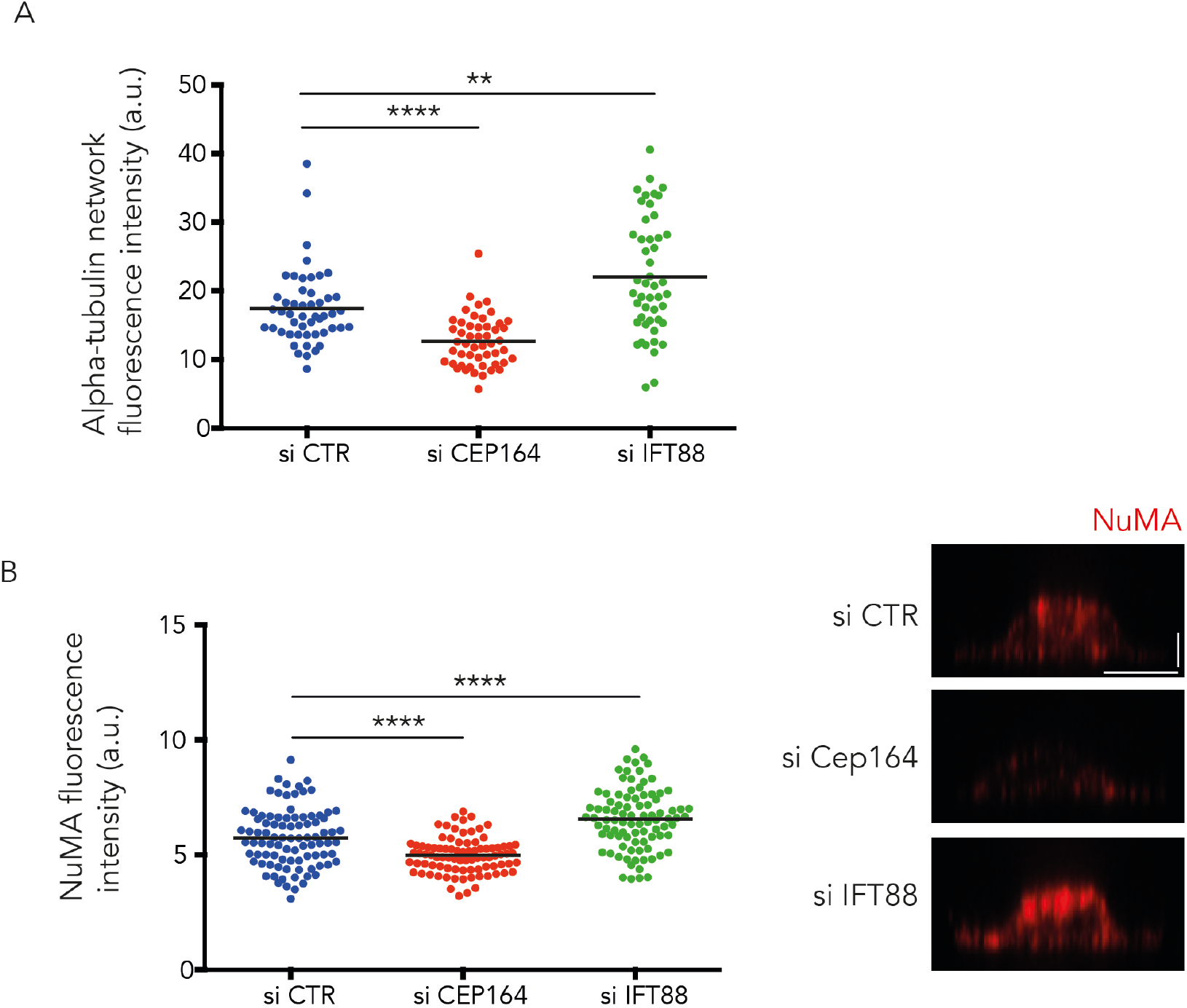
Implication of ciliogenesis effectors in microtubule network remodeling and apical NuMA recruitment. **A)** The graph shows the measurements of tubulin fluorescence intensity at the centrosome in RPE1 EGFP-centrin1 cells 4 hours post serum starvation (results of 2 independent experiments, n=100 cells per condition). **B)** The graph shows the measurements of NuMA fluorescence intensity at the centrosome in RPE1 EGFP-centrin1 cells 4 hours post serum starvation (results of 2 independent experiments, n=100 cells per condition). Images show representative side views of NuMA staining in response to siRNA treatments. XY scale bars represent 10 µm and Z scale bars represent 2,5 µm.

